# European scenarios for future biological invasions

**DOI:** 10.1101/2022.09.13.507777

**Authors:** Cristian Pérez-Granados, Bernd Lenzner, Marina Golivets, Wolf-Christian Saul, Jonathan M. Jeschke, Franz Essl, Garry D. Peterson, Lucas Rutting, Guillaume Latombe, Tim Adriaens, David C. Aldridge, Sven Bacher, Rubén Bernardo-Madrid, Lluís Brotons, François Díaz, Belinda Gallardo, Piero Genovesi, Pablo González-Moreno, Ingolf Kühn, Petra Kutleša, Brian Leung, Chunlong Liu, Konrad Pagitz, Teresa Pastor, Aníbal Pauchard, Wolfgang Rabitsch, Helen E. Roy, Peter Robertson, Hanno Seebens, Wojciech Solarz, Uwe Starfinger, Rob Tanner, Montserrat Vilà, Núria Roura-Pascual

## Abstract

1. Invasive alien species are one of the major threats to global biodiversity, ecosystem integrity, nature’s contribution to people and human health. While scenarios about potential future developments have been available for other global change drivers for quite some time, we largely lack an understanding of how biological invasions might unfold in the future across spatial scales.
2. Based on previous work on global invasion scenarios, we developed a workflow to downscale global scenarios to a regional and policy-relevant context. We applied this workflow at the European scale to create four European scenarios of biological invasions until 2050 that consider different environmental, socio-economic and socio-cultural trajectories, namely the European Alien Species Narratives (Eur-ASNs).
3. We compared the Eur-ASNs with their previously published global counterparts (Global-ASNs), assessing changes in 26 scenario variables. This assessment showed a high consistency between global and European scenarios in the logic and assumptions of the scenario variables. However, several discrepancies in scenario variable trends were detected that could be attributed to scale differences. This suggests that the workflow is able to capture scale-dependent differences across scenarios.
4. We also compared the Global- and Eur-ASNs with the widely used Global and European Shared Socioeconomic Pathways (SSPs), a set of scenarios developed in the context of climate change to capture different future socio-economic trends. Our comparison showed considerable divergences in the scenario space occupied by the different scenarios, with overall larger differences between the ASNs and SSPs than across scales (global vs. European) within the scenario initiatives.
5. Given the differences between the ASNs and SSPs, it seems that the SSPs do not adequately capture the scenario space relevant to understanding the complex future of biological invasions. This underlines the importance of developing independent, but complementary, scenarios focused on biological invasions. The downscaling workflow we presented and implemented here provides a tool to develop such scenarios across different regions and contexts. This is a major step towards an improved understanding of all major drivers of global change including biological invasions.

## INTRODUCTION

Invasive alien species (IAS) are species that have been introduced, established and spread beyond their natural range and have environmental or socio-economic impacts in the invaded area (CBD 2008; Blackburn et al. 2014, Bacher et al. 2018). IAS are recognized as one of the major threats to global biodiversity, regional economies, as well as human health and well-being (CBD 2008; Vilà et al. 2010; IPBES 2019; Novoa et al. 2021, Pysek et al. 2020). The numbers of IAS worldwide have steadily increased from 1950 onwards and will likely continue to accumulate in the future (Seebens et al. 2017, 2021). The projected increase in IAS numbers is mainly driven by increased global trade, socio-economic activities and climate change, among other environmental and societal drivers (Essl et al. 2020; Latombe et al. 2022). In conjunction with increased IAS numbers, it is expected that the impacts associated with IAS will also continue to grow (Pysek et al. 2020). Despite their prominent role in global biodiversity change (IPBES 2019), IAS are currently largely ignored in quantitative biodiversity projections (Lenzner et al. 2019), which highlights the need to comprehensively explore alternative future trajectories of biological invasions (Essl et al. 2019; Roura-Pascual et al. 2021).

To better understand how future biological invasions may unfold, several studies have explored the relationships between invasions and their individual drivers of global change, such as global trade (including wildlife trade; Seebens et al. 2015; Cardador et al. 2019; Sardain et al. 2019), climate change (Hellman et al. 2008; Hulme et al. 2017, Gallardo et al. 2017), land-use change (Decker et al. 2012; Walker et al. 2017) and human demography (Pyšek et al. 2010; Dawson et al. 2017). Moreover, only a few studies have assessed the relationship between invasions and multiple drivers simultaneously at the global scale (e.g. Latombe et al. 2022; Lopez et al. 2022). Importantly, drivers are not independent of each other, and due to the complex nature of biological invasions, predicting invasions requires the integrated assessments of a large set of drivers and their interactions in various socio-economic settings (Lenzner et al. 2019).

Scenario analysis is a well-established method to evaluate complex relationships among many drivers of change (e.g. Duinker & Greig 2007). Roura-Pascual et al. (2021) recently used scenario analysis to investigate the complex interactions between different global drivers of biological invasions, subsequently developing a set of global qualitative scenarios for biological invasions until 2050 (hereafter Global Alien Species Narratives; Global-ASNs). These scenarios describe potential alternative future social, political, socio-economic and environmental developments, with a special focus on biological invasions. Applying global scenarios to individual geographical regions is, however, challenging, as their underlying assumptions do not account for regional contexts (Chen et al. 2020, Bezerra et al. 2022, Latombe et al. in 2022). Therefore, downscaling global scenarios is essential to ensure that scenarios for specific regions incorporate both regionally important drivers and regional policy perspectives (Verburg et al. 2006; Chen et al. 2020). For example, Kok et al. (2019) recently adapted the global Shared Socio-economic Pathways (hereafter Global-SSPs; O’Neill et al. 2017) to the European context, which resulted in a set of four European shared socio-economic pathways (hereafter Eur-SSPs). The SSPs have been developed by the climate community to complement the initial climate change scenarios describing different levels of greenhouse gas emissions (i.e., the Representative Concentration Pathways; RCPs) and investigate the effect of different socio-economic trends on climate change (O’Neill et al. 2014; 2017). Given that the trajectories of biological invasions vary across spatial scales, it is important to develop and apply a protocol for downscaling global scenarios of biological invasions to finer resolutions. Europe is one of the regions where many biological invasions occur and counts with high density of data and research on alien species (Seebens & Kaplan 2022). Moreover, Europe has a long tradition regarding IAS regulation, but with research and management mostly at country level. Only relatively recently, efforts to integrate and coordinate transnational activities have been started (e.g. EU IAS Regulation 1143/2014). Thus, developing scenarios of biological invasions for Europe will help to identify research and knowledge needs, and some of the lessons learned might be useful for future exercises also in other regions.

Here, we develop and apply a participatory downscaling workflow to derive European scenarios (European Alien Species Narratives; Eur-ASNs) from the recently developed Global-ASNs (Roura-Pascual et al. 2021). We describe the downscaling process, reflect on the challenges we experienced, and provide guidance for future initiatives aimed at applying the ASNs to other regional contexts. In the next step, we present the resulting four Eur-ASNs for Europe until 2050 and highlight the novel information contained in them compared to Global-ASNs. Lastly, we quantitatively compare the Eur-ASNs with the Global-ASNs, Eur-SSPs and Global-SSPs to determine the extent to which the different scenarios align across scales (European vs. global) and scopes (ASN vs. SSP). The SSPs are frequently used to project biodiversity change (e.g., Di Marco et al. 2019; Leclère et al. 2020) despite the fact that their capacity to assess such change has been reported to be limited (Rosa et al. 2017; Pereira et al. 2020, Roura-Pascual et al. 2021). Therefore, our assessment will explore the possible differences between scenario initiatives, which is needed to understand their suitability for projecting future biological invasions.

## MATERIAL AND METHODS

### Developing European Alien Species Narratives (Eur-ASNs)

Future scenarios of biological invasions in Europe (i.e., Eur-ASNs) were developed during a workshop that was held online in two parts on 1–2 April 2020 and 30 September–2 October 2020 (see Roura-Pascual et al. 2022). A total of 35 participants from 12 European countries and three stakeholder groups (public administration, NGO/interested groups and academia) attended the workshop. Participants included invasion ecologists (23 participants), managers and policy-makers (8) as well as experts in global change and environmental history (3), and a specialist in scenario analyses (Supplementary Table S1). The Global-ASNs presented by Roura-Pascual et al. (2021) were used as the basis for downscaling the Eur-ASNs. Global-ASNs consisted of 16 individual scenarios, which were then grouped into four clusters based on their similarities and levels of invasion (see Box 1). The workshop participants were divided into four scenario breakout groups, each including representatives from all stakeholder groups. Each breakout group focused on one of four Global-ASN clusters, from which it selected one scenario to downscale to the European level (Figure 1). The selected scenarios were “Ruderal world” from cluster A, “Globalized corporation society” from cluster B, “Fairy tale” from cluster C and “Hipster/Techno society” from cluster D (for more information on main differences among the four selected scenarios, see Box 1; for specific details about scenario clusters, see Roura-Pascual et al. 2021). The breakout groups discussed which components of the scenario needed modification to compose a coherent scenario storyline for Europe from the present (using 2020 as baseline) until 2050. The participants focused on the same categories of socio-ecological drivers of biological invasions as the ones evaluated in Roura-Pascual et al. (2021) for the Global-ASNs: (i) political and institutional developments, (ii) socio-economic and demographic developments, (iii) culture, norms and values, (iv) technological development and science and (v) use of natural resources and ecological development. Additionally, a specific section in the Eur-ASNs was devoted to the dynamics and impacts of IAS at the European level. The four resulting Eur-ASNs cover a wide range of potential alternative future trends of social, political, socio-economic and environmental developments in Europe until 2050, with a special focus on biological invasions (Figure 2).

**Figure 1.**
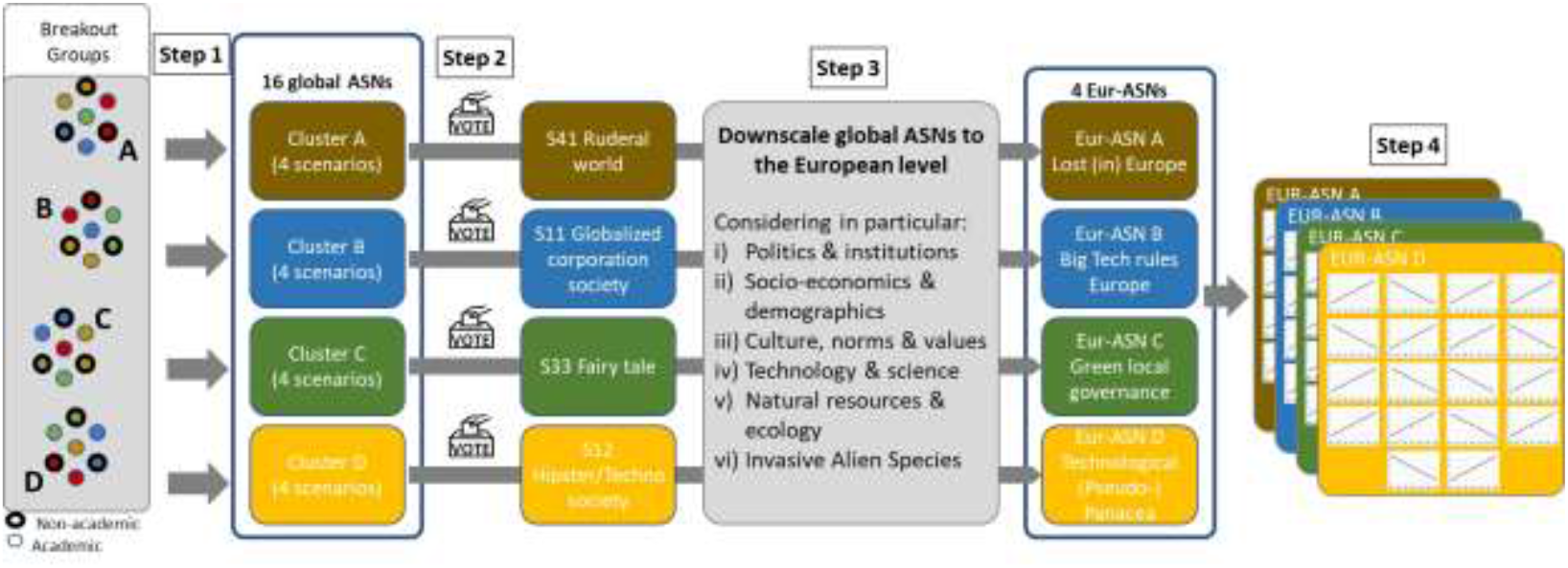
Conceptual description of the downscaling workflow to derive the Eur-ASNs from the Global-ASNs. Step 1: Workshop participants are assigned to four scenario breakout groups, each of which focuses on a different cluster of the global Alien Species Narratives (Global-ASNs, Roura-Pascual et al. 2021). Step 2: Each breakout group selects one Global-ASN from its assigned cluster. Step 3: Downscaling of the four selected Global-ASNs to the European context, resulting in four European Alien Species Narratives (Eur-ASNs). Step 4: Scoring the magnitude of change by 2050 for 26 variables widely used for quantification of socio-economic scenarios for each Eur-ASN.

**Figure 2.**
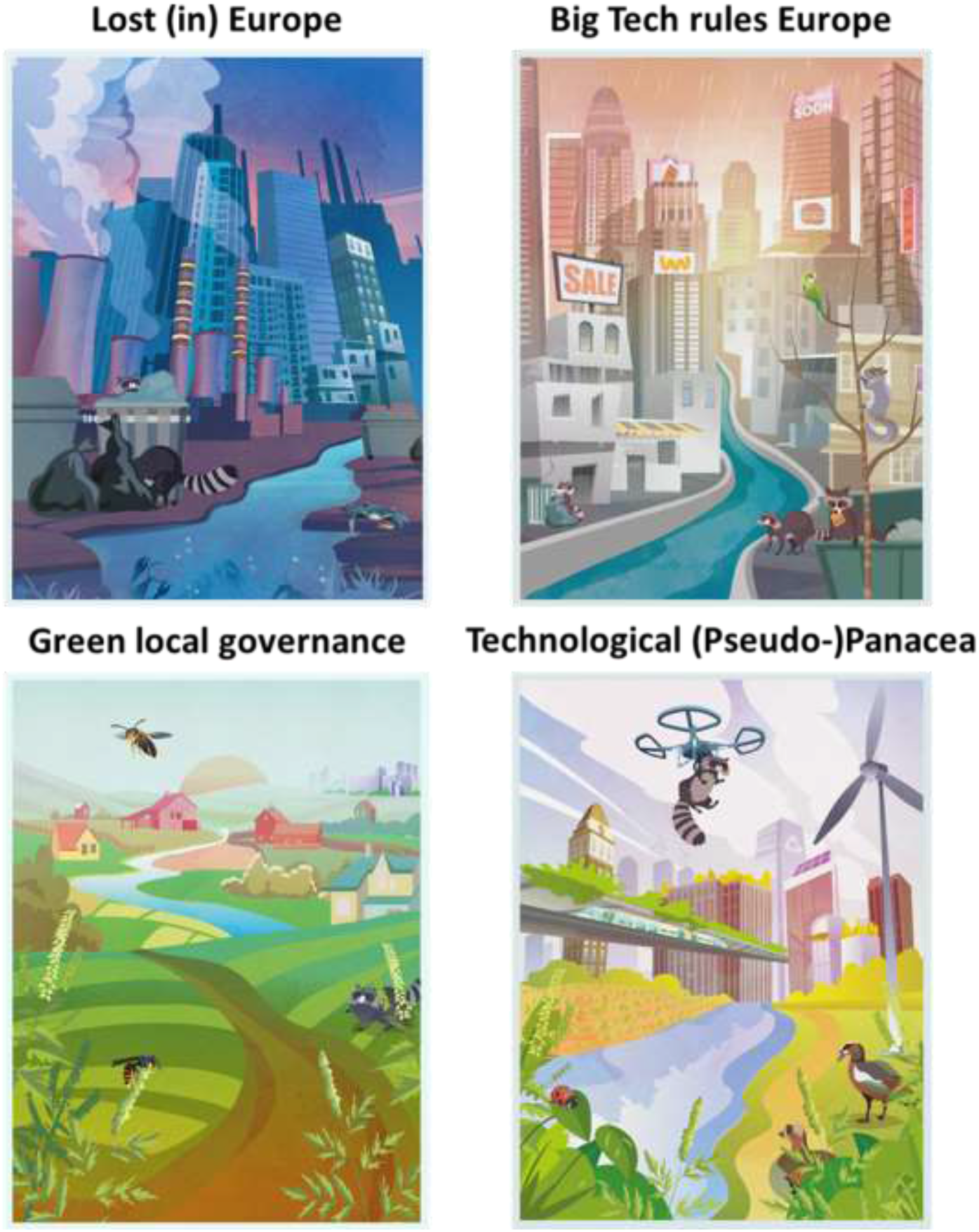
Artistic interpretations of the four European Alien Species Narratives depicting potential futures of Europe in 2050. Illustrations by K. Tsenova.

### Comparing Eur-ASNs with other scenarios

To enable comparison of the Eur-ASNs with Global-ASNs, Eur-SSPs and Global-SSPs, we reviewed the scientific literature to identify variables that were often used to quantify socio-economic scenarios (see consulted references in Supplementary Table S2). From the literature, we extracted 290 variables which were then assigned to 26 broad categories by (1) removing duplicates and resolving synonyms (e.g., land use and land-use change were grouped under “land use”) and (2) grouping together related variables (e.g., traditional biomass use, energy demand and energy supply were classified as “consumption and diet”). To facilitate variable interpretation and scoring, we assigned the 26 categories to one of the following six classes: (i) Demography, (ii) Economy & Lifestyle, (iii) Environment & Natural Resources, (iv) Human Development, (v) Technology and (vi) Policy & Institutions, providing a rationale for each variable (see Table S2).

Next, we used the 26 categories to characterize the 17 scenarios from the four different scenario initiatives (Eur-ASN: four scenarios, Global-ASN: four scenarios selected for downscaling, Eur-SSP: four scenarios, Global-SSP: five scenarios). Because the scenario Global-SSP2 has not been downscaled for Europe (Kok et al. 2019), we did not consider it in our analyses. For each scenario, we scored the anticipated change of each of the 26 categories on a 5-point Likert scale: strong increase, increase, no change, decrease, or strong decrease. The scoring reflected the assessment of a given variable change from 2020 to 2050 based on scenario storylines and, where available, scenario quantifications (see the complete list of consulted references in Table S2). The time horizon of 2050 was chosen to have a consistent end year for all scenarios covered by the different storylines. In most cases, scores reflected the absolute levels of change, but three categories (climate change, population growth and technological progress) were scored based on projected changes in their rate. For each scenario, three researchers from the scoring team (see Supplemental Table S1) independently assessed and then agreed on preliminary consensus scores. For transparency and transferability, those researchers provided a rationale for each assigned score. After that, the other four researchers from the scoring team reviewed and, if needed, revised preliminary consensus scores and rationales for each scenario. In the case of the Eur-ASNs, the workshop participants reviewed the consensus scores and rationales for the Eur-ASN they co-developed. Final scenario category scores and corresponding rationales are provided in Supplementary Table S2.

To quantitatively compare the scenarios, we analyzed the 17 scenarios and 26 variables using a non-linear principal component analysis (PCA) of the scenario variable scores from the expert assessment. We used the *ordPCA* function from the “ordPens” R package (version 1.0.0; Gertheiss & Hoshiyar 2021) with a shrinkage parameter λ = 0.001 to maximize the variance explained by the first two principal components (PC). The analysis was performed to understand how scenarios arranged along ordination axes and in relation to each other and which variables contributed most to this organization. Subsequently, we performed hierarchical clustering using a Euclidean distance measure within the two-dimensional space defined by the first two principal components (PCs) and the complete linkage algorithm as implemented in the *eclust* function from the “factoextra” R package (version 1.0.7; Kassambara & Mundt 2020). An optimal number of clusters was calculated using Elbow statistics implemented in the *fviz_nbclust* function in the “factoextra” R package (version 1.0.7; Kassambara & Mundt 2020). All analyses were performed in the R programming environment (version 4.1.1; R Core Team 2021).

## RESULTS

### European Alien Species Narratives (Eur-ASNs)

The four Eur-ASNs comprise comprehensive storylines developed during the workshop, describing different potential future trajectories of socio-economic development and biological invasions in Europe until 2050. Below, we present brief summaries of the storylines for each EUR-ASN, while the full narratives can be found in Supplementary Box S1 (additionally, see Figure 2 for illustrations of the Eur-ASN).

#### Eur-ASN A: Lost (in) Europe

(derived from Global-ASN A: “Ruderal World”)

By 2050, Europe has become increasingly isolationist, characterized by limited collaboration, international distrust and increasing heterogeneity in wealth among and within countries. The EU does not exist anymore and some former countries have even split up and formed smaller alliances. Political decisions are mainly driven by the egotism of those in power, but national governments are less powerful than today, whereas large corporations thrive, concentrate enormous economic power and strongly influence politics. Media are mainstreamed and biased, and there is only an apparent democracy with limited power of the general public. Essential services such as health care and education are no longer universal and access to them is mostly based on wealth and power. Scientific literacy is low overall and varies greatly among people. Scientific solutions and their societal uptake are driven by short-term thinking and economic interests, while problems that require long-term solutions are insufficiently addressed. Agriculture is based more on plant than animal production, with a long-term drop in productivity due to climate change impacts and shortages of imported fertilizer and energy. Biodiversity substantially degrades over time due to intensified agricultural land use to secure food production and due to climate change. EU biosecurity regulations dissolve with the disintegration of the EU and are replaced by minimal national regulations. The use of commercially important IAS for primary industries (e.g. forestry, aquaculture, mariculture) increases. While IAS introductions are lower than for a scenario with more international trade and tourism, the introductions are largely uncontrolled and investment in IAS management is low overall except for those IAS that directly threaten human livelihoods through impact on food production or human health.

#### Eur-ASN B: Big Tech Rules Europe

(derived from Global-ASN B: “Globalized corporation society”)

The recession of the 2020s and the stimulus response to bail out companies lead to a distrust in governments due to their failure to stop the crisis and to a desire of people to look after their own. This leads to companies having increased power over European policy and a lack of mechanisms to control them (and their ability to hide in tax havens and behind financial instruments) because they are multinational and difficult to halt, sanction, or regulate. People have economic power but are economically stressed and increasingly disconnected from the environment. Most Europeans focus on urban life and improving their situation in the city. The response to the crisis of the 2020s accelerates land abandonment and rural depopulation. Rural depopulation enables the widespread expansion of intensive and commercial farming for markets, often done by large companies. Citizens show little interest in nature, ecosystem services or biodiversity and have little knowledge about ecology and IAS. Substantial trade without biosecurity leads to an exponential increase in IAS. There is a large increase in IAS propagule pressure (due to intentional introductions and novel pathways of unintentional introductions) and a decrease in coordinated management. This leads to a further homogenization of IAS in urban areas and the appearance of substantial numbers of IAS in rural areas. Rural depopulation leads to a lack of knowledge and control of IAS in rural areas; in combination with increasing climate change this leads to disturbed ecosystems, in which IAS are difficult to control. European regulations to prevent the introduction of IAS exist, but the public sector has little power, thus most management actions to control IAS are implemented by companies. Control of IAS is focused on economically damaging IAS, but is often slow, scarcely coordinated and ineffective due to fragmented responses and lack of deeper knowledge due to limited collaboration and knowledge exchange. Trade and river transport increase the spread of IAS in freshwater and marine ecosystems.

#### Eur-ASN C: Green Local Governance

(derived from Global-ASN C: “Fairy Tale”)

By 2050, the EU as we know it (i.e. the common market promoting the exchange of goods, capital, services and labour) does not exist anymore and Europe sees the development of regionalism. It is not an increase in national populism and xenophobia, but a valorisation of local cultures and participatory democracy. Regional governments acquire greater influence because of the strong bottom-up participatory society, although there is still good cooperation among countries on certain political decisions or concerns (such as human health). Despite certain cooperation at a continental scale, there is a more superficial understanding of global (environmental) issues than currently. European society follows the degrowth paradigm, with less technological developments, and with the production of essential goods and services and consumption of local products. Eco-efficient, locally based production techniques are valued. People move from urban to rural areas. All these actions result in a reduction of GHG emissions and therefore a reduction of the impact of climate change. Remote working and remote learning is necessary and made easily accessible due to more distributed populations. Habitat fragmentation and increased land sharing are major pressures on the environment and the countryside, but mitigation measures, such as the creation of green corridors, are implemented. Nonetheless, some conservation efforts are less efficient due to the lack of a common environmental strategy. Because of isolation and reduced trade, the rate of introduction of new alien species coming from outside Europe decreases. However, the further spread of already established IAS is difficult to manage because of less efficient biosecurity measures, not coordinated at a continental scale.

#### Eur-ASN D: Technological (Pseudo-)Panacea

(derived from Global-ASN D: “Hipster/Techno Society”)

European nations in this scenario cooperate strongly, with fast technological advancement, large trade volumes and high biosecurity being the prime societal and policy objectives. Throughout Europe (beyond EU borders), agencies responsible for environmental policy are significantly strengthened and biosecurity legislation is strictly enforced. Despite strong cooperation, the reintroduction of intra-European border controls for biosecurity reasons leads to the end of free movement within the EU. European societies are highly urbanophile and concentrate in “Smart cities”. The urban lifestyle is supported by intensive, industrialized agriculture in rural areas and results in a decoupling from nature. Citizens strongly believe that technological progress can solve all current and future problems. Education is equally accessible to all parts of society and the general public acknowledges environmental issues (e.g. the impact of IAS). There is substantial cooperation regarding conservation and biosecurity measures, but policies are reactive rather than proactive. The strict regulations entail a degree of rigidity in societal life that fosters the development of rather complacent, passive societies. To a limited extent, implementation of the regulations varies among countries due to differences in types of governance, cultural legacies and values. Europe, and its individual nations, are engaged in an all-engulfing race to stay ahead of potential problems and of competitors by developing new technological solutions. Technological research and entrepreneurship is strongly promoted. Technologies for reducing the ecological footprint of various activities are available and implemented across Europe. Societies have a high, but not increasing ecological footprint. Catastrophic events, such as fires and floods, are under control using the latest technology, and further biodiversity loss is halted. Despite intensive global trade, the rate of IAS establishment and spread is low because of strong and diligent biosecurity measures, efficient risk assessments and other precautionary measures. As a result, different sets of IAS occur in cities and semi-urban environments as well as transportation networks outside cities and areas of intensive agriculture (i.e. predominantly in highly technological and artificial novel ecosystems). IAS management is supported by technological advances in automated and remote data collection with very high coverage, at large spatial and temporal scales and using standardized protocols in Europe.

### Comparison between Eur-ASNs and Global-ASNs

The PCA revealed that, as expected, the newly developed Eur-ASNs largely follow the logic of their global counterparts, with the respective pairs of Eur- and Global-ASNs clustering together in the PCA space (Figure 3). For certain variables, however, Eur-ASNs diverge from Global-ASNs (Figure 4). Between Eur-ASN A and Global-ASN A as well as between Eur-ASN B and Global-ASN B just one variable varied (“migration to/from Europe” and “population growth rate”, respectively, both varying from Increase to Decrease) (Figure 4). For Eur-ASN C and Global-ASN C, the variables “policy” and “inequality” switched from Strong increase and Strong decrease, respectively, at the global scale to No change at the European level. Lastly, Eur-ASN D and Global-ASN D showed the largest number of divergences, with four variables (“consumption and diet”, “mobility”, “climate change rate” and “social cohesion”) changing from Increase at the global scale to Decrease for Europe (Figure 4).

**Figure 3.**
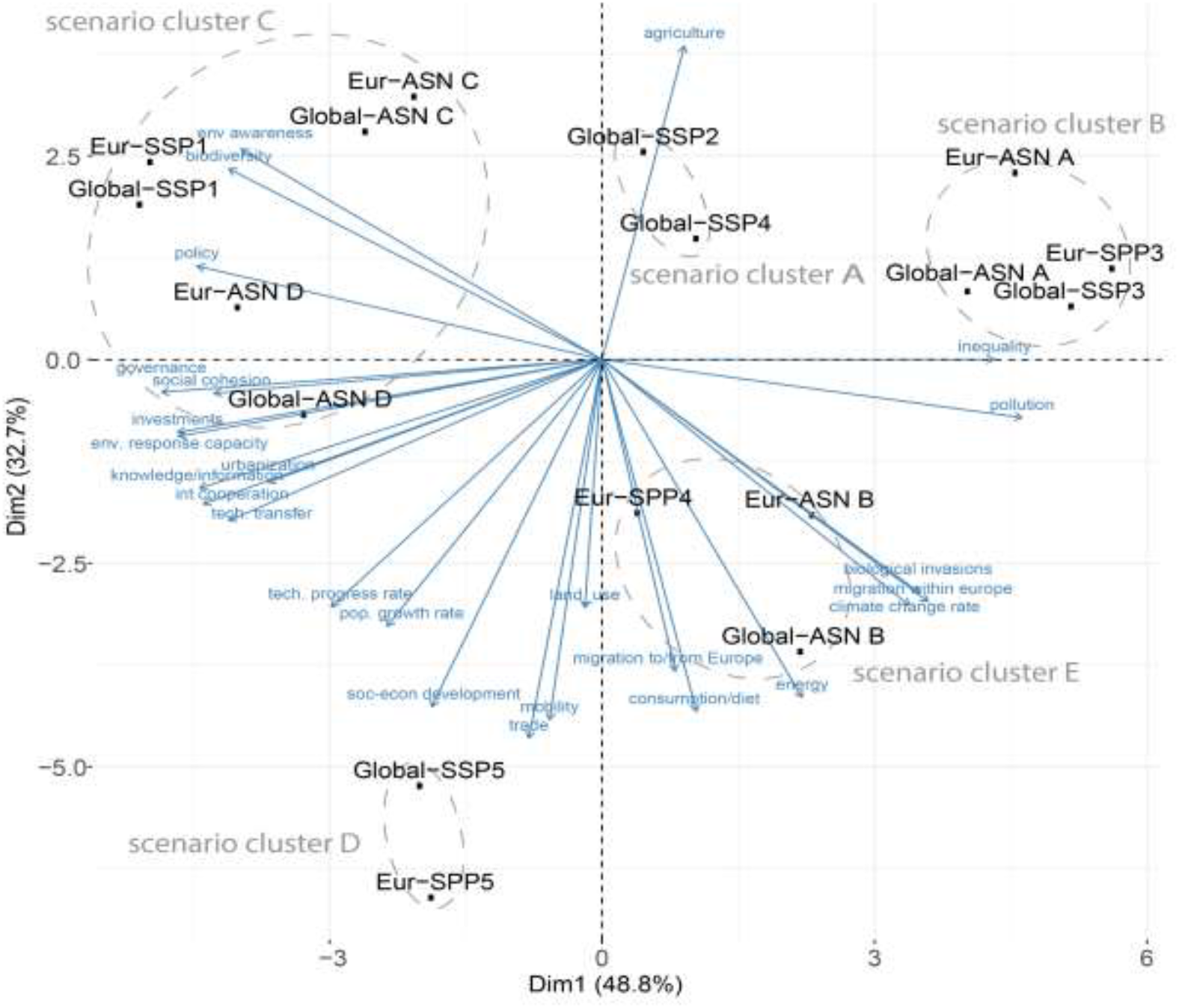
Biplot of the non-linear principal component analysis including the results from the hierarchical clustering. Optimal number of scenario clusters (indicated by dashed circles and indicated by “scenarios cluster” followed by a letter) was estimated using the Elbow method. Shown is the ordination of the European and Global ASNs and SSPs according to the first two principal components, explaining 81.5% of the variation. Arrows show factor scores for the scenario variables used in the analyses. Scenario abbreviations for the Alien Species Narratives (ASN) correspond to the following scenario names: Eur-ASN A = Lost (in) Europe; Eur-ASN B = Big Tech Rules Europe; Eur-ASN C = Green Local Governance; Eur-ASN D = Technological (Pseudo-)Panacea; Global-ASN A = Ruderal World; Global-ASN B = Globalized Corporation Society; Global-ASN C = Fairy Tale; Global-ASN D = Hipster/Techno Society.

**Figure 4.**
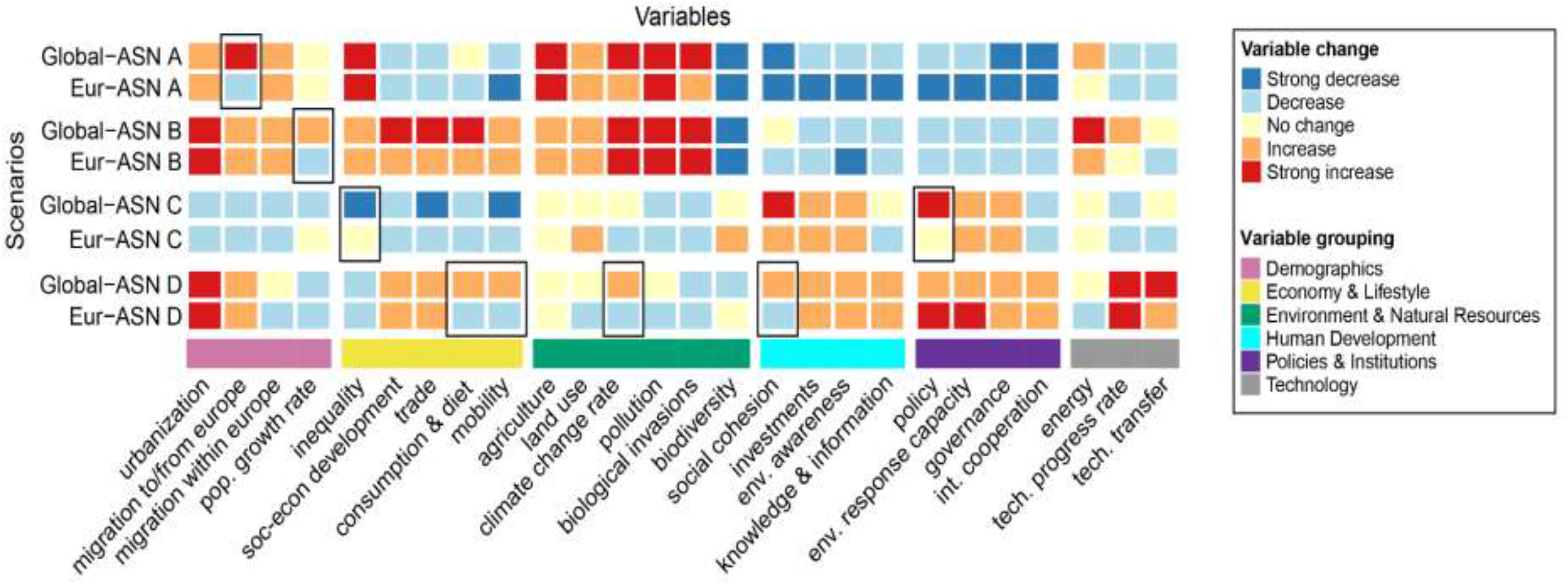
Comparison of projected future trends of 26 scenario variables between each newly developed European Alien Species Narrative (Eur-ASN) and its corresponding Global Alien Species Narrative (Global-ASN) based on expert scoring (from strong decrease to strong increase). Black frames highlight changes of a magnitude of at least two categories between scenarios, potentially resulting in reversed trends of scenario variables. The final consensus scores and rationales are provided in Supplementary Table S2. Scenario abbreviations correspond to the following scenario names: Eur-ASN A = Lost (in) Europe; Eur-ASN B = Big Tech Rules Europe; Eur-ASN C = Green Local Governance; Eur-ASN D = Technological (Pseudo-)Panacea; Global-ASN A = Ruderal World; Global-ASN B = Globalized Corporation Society; Global-ASN C = Fairy Tale; Global-ASN D = Hipster/Techno Society.

### Comparison across scenario initiatives and scales

The first two principal components (PCs) together captured 81.5% of variation (PC1 = 48.8% and PC2 = 32.7%; Figure 3). PC1 mainly captured information on policy and governance as well as inequality and pollution. PC2 was primarily related to migration, consumption and agriculture. The Elbow method estimated five clusters as optimal (i.e. the addition of more clusters or the reduction to less does not significantly increase the explained variation in the data) for classifying both the scenarios and the variables (Figures 3, 4).

#### Scenario clusters

Scenario cluster A includes Global-SSP 2 and Global-SSP 4, i.e. the two global SSPs that assume the continuation of historical trends or development towards a more unequal and divided society (Figures 3, 4). These trends include continuing and even intensifying increases in climate change and pollution, population growth, land use and urbanization as well as rising inequality, along with a decreasing interest in environmental policy and governance processes and international cooperation. Scenario cluster B includes Eur-SSP 3, Global-SSP 3, Eur-ASN A and Global-ASN A (Figures 3, 4). All scenarios in this cluster assume an increasingly isolationist world that strongly hampers international cooperation in policy, trade and transport. Consequently, social inequalities increase and environmental issues are only tackled at the national scale, if at all, leading to higher pollution, climate change and biodiversity loss. Scenario cluster C is centered around the premises of sustainable development, changes in consumption as well as reactive technological societies and comprises Eur-ASN D, Global-ASN D, Eur-SSP 1, Global-SSP 1, Eur-SSP 3, Eur-ASN C and Global-ASN C (Figures 3, 4). These scenarios assume strong confidence in governments and public actors, resulting in improved policy, governance and hence societal cohesion, although at different spatial scales. At the same time, there is a strong valuation of nature and the environment, leading to increased biodiversity and reduced climate change, pollution and land conversion. Notably, while the Global-ASN and Eur-ASN C & D are in the same cluster they differ in terms of urbanization, trade, consumption and technological development (Figures 3, 4). Scenario cluster D includes Eur-SSP 5 and Global-SSP 5 (Figures 3, 4), which focus strongly on fossil fuel-driven development in a highly connected world that is mainly oriented towards markets and large businesses. While most variables are projected to follow a positive trend, inequality, biodiversity and environmental awareness are expected to strongly decrease (Fig. 5). Finally, scenario cluster E focuses on market-oriented technology and includes scenarios Eur-ASN B, Global-ASN B and Eur-SSP 4 (Figures 3, 4). All these scenarios represent an internationally well-connected world driven by international corporations and business elites, with strong economies and weak governments.

**Figure 5.**
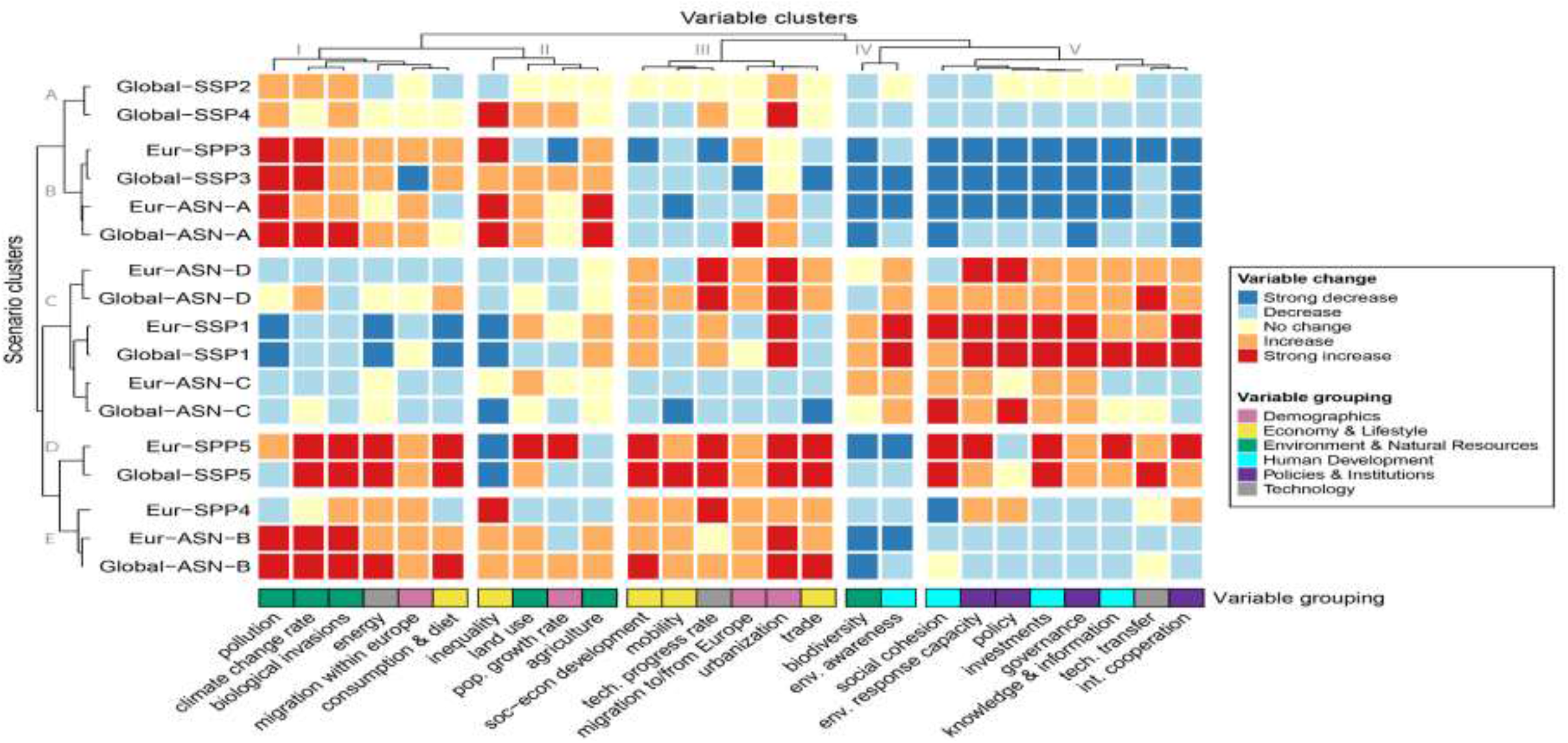
Comparison among the scenario initiatives (Global-SSP, Eur-SSP, Global-ASN, Eur-ASN) based on the variable scoring by the expert group. Clustering of scenarios and variables is based on the results of a non-linear principal component analysis using the Elbow method to determine the optimal number of clusters, which are represented with capital letters in Y-axis. Scenario abbreviations for the Alien Species Narratives (ASN) correspond to the following scenario names: Eur-ASN A = Lost (in) Europe; Eur-ASN B = Big Tech Rules Europe; Eur-ASN C = Green Local Governance; Eur-ASN D = Technological (Pseudo-)Panacea; Global-ASN A = Ruderal World; Global-ASN B = Globalized Corporation Society; Global-ASN C = Fairy Tale; Global-ASN D = Hipster/Techno Society. Scenario abbreviations for the European and Global SSPs correspond to the original ones published in O’Neill et al. (2017) and Kok et al. (2019).

This development is very resource-intensive, resulting in high rates of climate change, consumption and pollution leading to strong negative effects on biodiversity.

#### Variable groups

The clustering algorithm grouped variables (Figure 5, upper margin) by pollution, climate change rate, biological invasions, energy use, migration within Europe and consumption & diet (group I); social inequality, land-use change, population growth rate and agriculture, (group II); socio-economic development, mobility, technological progress rate, migration to/from Europe, urbanization and trade (group III); biodiversity and environmental awareness (group IV); and societal cohesion, environmental response capacity, policy, investments, governance, knowledge & information, technology transfer and international cooperation (group V).

Scenario clusters A and B are characterized by quite similar changes in their variable values (Figure 5). They show mainly an increase in variables of groups I and II and a decrease in variables of groups IV and V. Differences mainly lie in variable group III, where scenario cluster A shows no change for most of the considered variables while scenario cluster B decreases. In contrast, scenario clusters D and E both show increase in variables from groups I and III and decrease in variable group IV. They mainly diverge in groups II and V. Finally, scenario cluster C shows decreasing trends for variable groups I and II and an increasing trend for variable group IV, in contrast to other scenario clusters. Interestingly, this is the only cluster where biological invasions show a decreasing trend. Scenario clusters C and D show increasing trends for variables from group V, while the other scenario clusters show a mainly decreasing trend in these variables.

## DISCUSSION

In this study, we introduce the first European scenarios for future biological invasions, which were obtained by downscaling the Global-ASNs (Roura-Pascual et al. 2021). Overall, the European scenarios aligned well with their global counterparts but partially diverged from the scenario space covered by the SSPs. This may suggest that the ASNs are better at capturing biodiversity changes in general and changes in biological invasions in particular. This finding is consistent with other calls for independent biodiversity scenarios to replace the frequently used SSPs, originally designed by the climate community (Pereira et al. 2020).

### Comparing global and European scenarios of biological invasions

The Eur-ASNs cover a wide range of socio-economic, environmental and socio-cultural variables and anticipate substantially different levels of invasion across Europe, ranging from low decrease (Eur-ASN C) to no change (Eur-ASN D) and substantial increase (Eur-ASN A and Eur-ASN B). We found overall good alignment between the biological invasion scenarios across spatial scales. This was to be expected, given that the Eur-ASNs were inferred directly from their global counterparts (Figure 1). Nonetheless, for a few variables, the trends in the Eur-ASNs substantially deviate from those at the global scale. Specifically, unlike their global counterparts, certain Eur-ASNs assume a decrease in migration to/from Europe, population growth rate, consumption and diet, mobility, climate change rate, social cohesion and policy. Identifying contextual differences across spatial scales is a core aim of any downscaling approach. Such deviations show the relevance of context-dependent patterns on biological invasions, where global drivers might interact differently at smaller scales (González-Moreno et al. 2013). These findings agree with the results obtained by Kok et al. (2018), who found the Eur-SSPs to be well-aligned with the Global-SSPs, besides the existence of some deviations between scenarios.

For the Eur- and Global-ASN A (“Lost (in) Europe”, derived from “Ruderal World”), we have one such notable deviation for migration to/from Europe. These scenarios relate to an increasingly isolationist world, with high potential for rising inequalities and risks of conflicts likely resulting in increased migration worldwide. At the European scale, however, increasingly isolationist tendencies and hostility towards non-European migrants, together with the reintroduction of intra-European borders and the abolishment of the Schengen agreement proposed for Eur-ASN A, will likely reduce migration within and to Europe.

In Eur- and Global-ASN B (“Big Tech Rules Europe”, derived from “Globalized corporation society”), trends for population growth rate are reversed across the scales. At the global scale, globalization and socio-economic development is assumed to continue, likely resulting in future population growth rates similar to the currently observed (Lutz et al. 2018). At the European scale, however, the population growth rate will likely decline due to a continuation of the aging of the population and decreasing birth rates particularly in rural areas (MacDonald et al. 2000; Lutz et al. 2018).

Eur- and Global-ASN C (“Green Local Governance”, derived from “Fairy Tale”) show opposite trends for inequality and policy. At the global level, societies are assumed to be highly democratic, self-sufficient and egalitarian, with reduced conflict potential. Consequently, inequality will decline. Within Europe, however, inequalities will slightly increase, given that different starting resources like technology and infrastructure will not be readily redistributed among countries due to political regionalization. At the same time, the scenario assumes a movement from urban to rural areas that may reduce inequality at the European level. Policy, however, becomes a priority, with a primer on bioconservatism and biosecurity resulting in effective policies at local and regional scales. At the European scale, the disintegration of the EU and the increased regionalization will likely complicate efficient policy measures and policy efforts will be undermined by countries with the weakest policies.

Finally, Eur- and Global ASN D (“Technological (Pseudo-)Panacea”, derived from “Hipster/Techno Society”) show most deviations across scale, including consumption and diet, mobility, climate change rate and social cohesion. Globally, a high but stabilized ecological footprint of future societies owing to a moderate increase in consumption and a shift towards less impactful diets. Given the globally high and increasing standard of living and associated high ecological footprints climate change rate will keep increasing. The high standard of living with strong democratic societies overall translates into increasing social cohesion and paired with environmental awareness and technological advancements people will likely become more mobile globally. At the European scale, technological development will increase production efficiency and the promotion of eco-friendly products will lead to declining trends in consumption. Technological advances and innovation will also reduce the climate change rate due to advancements in climate engineering, renewable energy production, low-carbon technology development and increased energy efficiency. Social cohesion at the European level is expected to decrease following the loss of important cultural aspects of human and non-human lives. Societies become more individualist and a vibrant cultural sector is of low importance. Finally, mobility will likely decrease due to intra-European border controls and higher environmental awareness hampering the motivation to travel.

Overall, all Eur-ASNs result in different levels of biological invasions, neglecting potential futures where biological invasions can be effectively mitigated and reduced. This trend might be influenced by the selection procedure for developing the Eur-ASNs based on the Global-ASNs, which have the same shortcomings. Future work should develop more optimistic scenarios of biological invasions to outline pathways towards a desirable future. Such future efforts could, for example, utilize the Nature Futures Framework that aims towards positive scenarios for biodiversity (Pereira et al. 2020).

### Comparing the ASNs with the SSPs

We found that all Eur-ASNs except Eur-ASN B were well aligned with both the Global-ASNs and Eur-SSPs. Similarly, Roura-Pascual et al. (2021) found differences between ASNs and SSPs at the global scale. Interestingly, Eur-ASNs and Eur-SSPs match better than their global counterparts. This convergence among scenarios designed for different purposes suggests that using the appropriate scale is important to generate comparable scenarios. At the same time, some of the scenario space occupied by the SSPs is not covered by the recently developed Eur-ASNs and the Global-ASN considered in this exercise. Specifically, Global-SSPs 2 & 4 and Global and Eur-SSP 5 represented two separate clusters (clusters A and D, respectively, Figure 3).

The mismatch between ASNs and SSPs highlights that different scenario initiatives are necessary to explore different facets of environmental change. The SSPs, together with the future climate change scenarios (i.e., the Representative Concentration Pathways; RCPs), are frequently used to project biodiversity change (e.g., Leclère et al. 2020; Pereira et al. 2020). However, the modeling community, as well as decision-makers and practitioners, need to be aware that such models and scenarios are climate-centric and hence limited in their assessment of biodiversity change (Rosa et al. 2017). Hence, comparisons of the widely used SSPs with biodiversity-centric scenarios like the ASNs are crucial to identify biodiversity-specific aspects. The Nature Futures Framework, developed by IPBES (Pereira et al. 2020), facilitates scenario development with a special focus on positive scenarios of biodiversity change, integrating different value systems. Exploring synergies between the scenarios developed by the Nature Futures Framework community and the biological invasion scenarios has large potential to understand future synergistic effects at the biodiversity-health-society interface and to develop mitigation and adaptation strategies. Further efforts would benefit from broader expertise and consider the local perspective provided by a larger variety of disciplines and key stakeholders, including government and local and indigenous communities.

### Downscaling workflow from global to regional scenarios of biological invasions

The downscaling workflow was based on a participatory process involving experts with diverse scientific and non-scientific backgrounds (Roura-Pascual et al. 2022). The development of the Eur-ASNs and the 26-category scoring may be sensitive to the expertise involved. However, due to the large number of experts involved and that they reached a consensus on key regional drivers of biological invasions at the European scale, we believe that the overall credibility of the Eur-ASNs should be high (Sutherland and Burgman 2015). High diversity of expertise, perspectives and disciplines among the participants is crucial to ensure that all environmental, socio-economic, political and societal facets relevant to the regional context are well represented (Hannagan and Larimer 2010; Krueger et al. 2012; IPBES 2016). The downscaling workflow used in this exercise can be adopted in other contexts, for example to downscale the biological invasions scenarios to other geographical regions or specific taxonomic groups. Therefore, we strongly encourage others to perform similar studies to increase the variety of potential future scenarios for biological invasions.

### Conclusions

The Eur-ASNs developed here cover a wide range of plausible futures, which can contribute to assessing the future of biological invasions in Europe and provide an important step to address one of the major drivers of biodiversity loss. The identification of plausible futures for biological invasions in Europe, together with assessment of key drivers to reduce or increase the level of invasions, can be used as a starting point for developing quantitative scenario projections and improving management strategies of biological invasions (Roura-Pascual et al. 2022). Such management should aim to establish action plans flexible enough to be adapted to various future developments. Extending our approach and scenarios to other geographical areas and time horizons would increase our understanding of how biological invasions may unfold across space and time by accounting for environmental, socio-economic and societal trends. We are conscious that the level of information on Europe may not be available in other regions, but we hope that the lessons learned here will help identify key information and knowledge gaps for other regions and contribute to perform future scenarios exercises in regions under-represented in biological invasion research (Núñez and Pauchard 2010).

## Author contribution statement

**Cristian Pérez-Granados, Bernd Lenzner, Marina Golivets, Wolf-Christian Saul:** Conceptualization, Methodology, Formal analysis; Investigation, Writing – Original Draft, Writing – Review and Editing, Visualization. **Jonathan M. Jeschke, Franz Essl, Núria Roura Pascual**: Conceptualization, Methodology, Investigation, Writing – Review and Editing, Visualization, Project administration. **Garry D. Peterson, Lucas Rutting, Guillaume Latombe**: Conceptualization, Investigation – Workshop and Online Contribution, Writing – Review and Editing. **Tim Adriaens, David Aldridge, Sven Bacher, Rubén Bernardo-Madrid, Lluís Brotons, Belinda Gallardo, Piero Genovesi, Pablo González-Moreno, Ingolf Kühn, Petra Kutleša, Chunlong Liu, Konrad Pagitz, Teresa Pastor, Wolfgang Rabitsch, Helen E. Roy, Peter Robertson, Hanno Seebens, Wojciech Solarz, Uwe Starfinger, Rob Tanner, Montserrat Vilà:** Investigation – Workshop and Online Contribution, Writing – Review and Editing. **Brian Leung, Aníbal Pauchard:** Investigation – Online Contribution, Writing – Review and Editing.

## Acknowledgements

This research was funded through the 2017-2018 Belmont Forum—BiodivERsA international joint call for research proposals, under the BiodivScen ERA-Net COFUND program, through the AlienScenarios (https://alien-scenarios.org/) and InvasiBES (http://elabs.ebd.csic.es/web/invasibes) projects, with the following funding organizations: Spanish State Research Agency (MCI/AEI/FEDER, UE, PCI2018-092939; MV; PCI2018-092986; BG; PCI2018-092966; CPG, NRP), German Federal Ministry of Education and Research (BMBF; grants 16LC1807A, 16LC1807B and 16LC1807C; HS, W-CS, JMJ, MG and IK), Austrian Science Foundation (FWF; I 4011-B32; BL and FE), Swiss National Science Foundation (SNSF; 31BD30_184114; SB). We would like to acknowledge Spyridon Flevaris, Marcus Hall and Jörg Priess for their helpful contribution during the workshops and scenario development. We also thank Kris Tsenova for creating the scenario illustrations. CPG acknowledges the support from Ministerio de Educación y Formación Profesional through the Beatriz Galindo Fellowship (Beatriz Galindo – Convocatoria 2020). AP was funded by ANID/BASAL FB210006.

## Boxes

### Box 1

**Global Alien Species Narratives (ASNs) based on Roura-Pascual et al. (2021)**

There are a total of 16 Global-ASNs grouped into four clusters introduced by Roura-Pascual et al. (2021). Clusters A and B correspond to futures with high levels of biological invasion, while clusters C and D are characterized by low or medium invasion levels (see Roura-Pascual et al. 2021 for full descriptions and methodology).

- Scenarios in cluster A (including “Ruderal world”) are characterized by an irresponsible and inefficient use of natural resources with high reliance on fossil fuels; international trade decelerates but without reduction in the rate of climate change or introductions of IAS.
- Cluster B comprises scenarios (including “Globalized corporation society”) in which the political agendas are driven by global economic interests, with high productivity and international trade responsible for high consumption of natural resources; an urban life-style dominates, environmental concerns receive little attention and the levels of biological invasions are high.
- Cluster C is dominated by scenarios (including “Fairy tale”) with high levels of social and environmental awareness and regulation, which is implemented regionally but incorporates global environmental needs; global trade and human footprint are reduced and the levels of invasions are low. There is a tendency towards the development of environmental regulations at the regional scale, which hampers their effectiveness in the absence of effective higher level coordination. Nevertheless, regional levels of invasions are low compared to the 2020 level.
- Finally, Cluster D includes scenarios (including “Hipster/Techno society”) that are characterized by a good balance between regional and global governance systems, as well as high availability and utilization of novel technologies that provide solutions to mitigate the impact of some environmental pressures; invasion levels are intermediate because even though global trade is elevated, effective biosecurity programs are widely implemented.

