## Supplementary material for "European scenarios for future biological invasions"

**Table S1.** Workshop participants involved in the development of the four European Alien Scenarios Narratives, indicating their country and expertise. (*) indicates members of the workshop facilitation team, (^) indicates members of the scoring team.

| **Affiliation country** | **Stakeholder type** | **Stakeholder expertise** | **Name of participant** |
|---|---|---|---|
| Austria | Academia | Invasion ecology | Franz Essl^ |
| Austria | Academia | Invasion ecology | Bernd Lenzner^ |
| Austria | Academia | Policy and management | Konrad Pagitz |
| Austria | Public administration | Invasion ecology | Wolfgang Rabitsch |
| Belgium | Academia | Invasion ecology | Tim Adriaens |
| Belgium | Public administration | Policy and management | Spyridon Flevaris |
| Croatia | Public administration | Policy and management | Petra Kutlesa |
| China | Academia | Invasion ecology | Chunlong Liu |
| France | Public administration | Policy and management | François Diaz |
| France | Public administration | Policy and management | Rob Tanner |
| Germany | Academia | Invasion ecology | Marina Golivets^ |
| Germany | Academia | Invasion ecology | Jonathan M. Jeschke*^ |
| Germany | Academia | Invasion ecology | Ingolf Kühn |
| Germany | Academia | Invasion ecology | Jörg Priess |
| Germany | Academia | Invasion ecology | Wolf-Christian Saul*^ |
| Germany | Academia | Invasion ecology | Hanno Seebens |
| Germany | Academia | Invasion ecology | Uwe Starfinger |
| Italy | NGO / Interest group | Policy and management | Piero Genovesi |
| Netherlands | Academia | Scenario analysis | Lucas Rutting* |
| Poland | Public administration | Policy and management | Wojciech Solarz |
| Spain | Academia | Global change | Lluís Brotons |
| Spain | Academia | Invasion ecology | Rubén Bernardo |
| Spain | Academia | Invasion ecology | Belinda Gallardo |
| Spain | Academia | Invasion ecology | Cristian Pérez-Granados*^ |
| Spain | Academia | Invasion ecology | Núria Roura-Pascual*^ |

| Affiliation country | Stakeholder type | Stakeholder expertise | Name of participant |
|---|---|---|---|
| Spain | Academia | Invasion ecology | Montserrat Vilà |
| Spain | NGO / Interest group | Policy and management | Teresa Pastor |
| Sweden | Academia | Sustainability Science | Garry D. Peterson* |
| Switzerland | Academia | Invasion ecology | Sven Bacher |
| Switzerland | Academia | Environmental history | Marcus Hall |
| United Kingdom | Academia | Invasion ecology | David Aldridge |
| United Kingdom | Academia | Invasion ecology | Guillaume Latombe |
| United Kingdom | Academia | Invasion ecology | Peter Robertson |
| United Kingdom | Academia | Invasion ecology | Helen E. Roy |
| United Kingdom | NGO / Interest group | Invasion ecology | Pablo González-Moreno |

**Table S2.** Name, rationale, consensus score and consensus justification for each of the 26 variables considered to characterize the scenarios of all four scenario initiatives (Eur-ASN: 4 scenarios, Global-ASN: 4 selected scenarios, Eur-SSP: 4 scenarios, Global-SSP: 5 scenarios).

https://figshare.com/articles/dataset/Supplementary_Table_S2/20776207

**Box S1**. Detailed description of scenarios for biological invasions in Europe …

**SCENARIO Eur-ASN A: LOST (IN) EUROPE**

| | |
|---|---|
| **Scenario breakout group** | Tim Adriaens, François Diaz, Franz Essl, Pablo González-Moreno, Jonathan Jeschke (facilitator), Ingolf Kühn, Helen Roy, Hanno Seebens, Uwe Starfinger |
| **Scenario key features** | **Keywords:** isolationist Europe, disintegration of the EU, loss of European identity, rise of populism, concentration of political power, international trade reduction, science illiteracy, increased land use, environmental degradation<br>**IAS specifics:** loss of EU regulations due to EU disintegration, rise of commercial IAS for primary industries (e.g. forestry, aquaculture, mariculture), largely uncontrolled IAS introductions, overall low IAS management (except those IAS that directly threaten human livelihoods), increased IAS impacts and spread due to climate change and increased land use |
| **Scenario downscaled from** | Global-ASN S41 'Ruderal World' (Scenarios Cluster A, see Roura-Pascual et al. 2021) served as the basis for downscaling to the European level |

### SCENARIO SUMMARY

By 2050, Europe has become increasingly isolationist, characterized by limited collaboration, international distrust and increasing heterogeneity in wealth among and within countries. The EU does not exist anymore and some former countries have even split up and formed smaller alliances. Political decisions are mainly driven by the egotism of those in power, but national governments are less powerful than today, whereas large corporations thrive, concentrate enormous economic power and strongly influence politics. Media are mainstreamed and biased, and there is only an apparent democracy with limited power of the general public. Essential services such as health care and education are no longer universal and access to them is mostly based on wealth and power. Scientific literacy is low overall and varies greatly among people. Scientific solutions and their societal uptake are driven by short-term thinking and economic interests, while problems that require long-term solutions are insufficiently addressed. Agriculture is based more on plant than animal production, with a long-term drop in productivity due to climate change impacts and shortages of imported fertilizer and energy. Biodiversity substantially degrades over time due to intensified agricultural land use to secure food production and due to climate change. EU biosecurity regulations dissolve with the disintegration of the EU and are replaced by minimal national regulations. The use of commercially important IAS for primary industries (e.g. forestry, aquaculture, mariculture) increases. While IAS introductions are lower than for a scenario with more international trade and tourism, the introductions are largely uncontrolled and investment in IAS management is low overall except for those IAS that directly threaten human livelihoods through impact on food production or human health.

### IN DETAIL: WHAT DOES THIS SCENARIO IMPLY FOR EUROPE UP TO 2050?

#### REGARDING BROAD, CONTEXTUAL DEVELOPMENTS

**1. Political and institutional developments**

Rising pressures over time increasingly lead to an isolationist world, little collaboration and international distrust. Political instability leads to several changes of European borders (due to segregationist movements) and risks of wars and conflict over natural resources and essential goods (e.g. medical supplies, food, natural resources, energy, fossil fuels). The EU does not exist anymore, and tensions within countries result in many small independent regions. On the other hand, regional alliances between countries become more prominent, mostly based on national socio-economic interests and regional market mechanisms or common protective policies towards human migration, border protection, internal markets and food security (e.g. Scandinavian and Mediterranean countries flock together; Germany and Austria strongly work together, the Benelux is reinforced). Alliances are, however, based on mutualistic benefits to national societies rather than solidarity. Policy is short-sighted and egotistic (e.g. politicians are strongly influenced by lobbies and big corporations), leading towards a less humanitarian society with reduced solidarity. Governance moves towards centralization, and individual leaders are limiting regional governments' competences. Multinational corporations profit from loosened government control and dominate markets, enforcing oligopolies and monopolies. Similarly, media are mainstreamed and politically biased. There has been a rise of demagogues and national parties, and although there is an apparent democracy, public participation in decision making is rather limited.

**2. Socio-economic and demographic developments**

Inequalities among and within countries increase. Some regions in Europe are seeing their economies grow, but others, such as large parts of the West and the Southeast Mediterranean, experience economic decline. Thus, there are big disparities in per-capita income across the EU. Essential services such as health care are not universal anymore but mostly based on wealth and power. Most people will receive relatively poor education, whereas the elite will go to costly private schools, colleges and universities. The socio-economic inequalities cause large human migration pressure. Changes in diets and/or malnutrition become regionally endemic. Disease increasingly contributes to human mortality. Further, there is an increasing heterogeneity among regional clusters in demographic patterns. Although at continental level the human population remains stable, some countries with a more ruderal economy will have a higher population growth based on higher birth rate. Average life expectancy is decreasing: while rich people are long-lived, others have a lower life expectancy than today. Tourism becomes more regional and limited, restricting the movement of people and goods.

**3. Culture, norms and values**

Self-focused strategies, populist movements, xenophobia and radicalism dominate, including hostility towards migrants and refugees, and the emergence of currently unseen levels of violence and conflict. Long-term environmental issues (e.g. climate change, nitrogen deposition, soil erosion, lowering groundwater levels, wildlife extinction) are considered irrelevant. There is a move from a European identity towards more extreme national and regional identities based on language and common history. Typical aspects of European identity (inclusiveness, cosmopolitanism, tolerance, transnationalism, privacy, freedom of speech) are lost. People are less interested in learning about other cultures, their history and languages, and this is reflected in educational curricula which focus more on national identity. Grass-root movements are still active, including movements on food production (e.g. permaculture), drinkable water for poor people and climate-social justice movements. However, these movements are marginal, subversive, vocal minorities that are mostly silenced, festering frustratedly and largely hidden, e.g. in the dark web.

**4. Technological developments, science**

Scientific literacy is low overall, but with huge differences among different groups of people. Scientific solutions and their societal uptake are driven by short- and medium-term challenges, while long-term questions are insufficiently addressed. Innovation in general is low to moderate. Investments in public research and development are generally low and based on self-interest of major stakeholders. Individual companies (particularly the global players) will, however, invest much money into research and development; their findings are not publicly shared, giving the companies even more power. Some of these innovations will help solve IAS problems that threaten critical sectors. There is a tendency toward elite science, and countries are investing in scientific and innovative niches and high-profile prestige projects that benefit the few more than the general good or the public interest. Employment for ecologists is virtually non-existent, and scientific curricula are directed towards food production, pest and disease control. Soft sciences (particularly social sciences and philosophy) are little valued and practiced.

**5. Natural resources, ecological developments**

In the short/medium term, there are moderate impacts on biodiversity, but over time strong degradation of biodiversity occurs due to aggressive expansion of land use to secure food production, and due to climate change. Agriculture is based more on plant than on animal production, with a long-term drop in productivity due to climate change impacts, and a lack of imported fertilizers and oil. As resources are limited and global trade is low, there is a dietary shift toward a more plant-based diet, including nuts, grains and legumes; meat production decreases and agriculture is following this trend, trying to maximize plant production per surface area. Landscape heterogeneity is decreasing, with harder borders between natural areas and agricultural land. Little surface area is left unmanaged, and the pressure to use land for agricultural production is huge. Water will, like oil, become a scarce resource for several countries due to climate change, leading towards treating water as a key commodity.

### REGARDING SPECIFIC IAS DYNAMICS AND MANAGEMENT

Continental biosecurity regulations in plant health, animal health and biodiversity regimes are mostly downscaled to national levels, with little collaboration across borders. There is no cross-sectoral approach towards the conservation and sustainable use of natural resources, the preservation of biodiversity or the importance of the IAS problem. National nature conservation legislation is not enforced anymore. The EU ceases to exist in 2030, thus all EU regulations do not exist anymore by then. The Bird and Habitat Directive, as well as the Flood Directive, the Water Framework Directive and the Marine Strategy Framework Directive do not survive a fitness check and are abandoned. The Ballast Water Convention is loosened too. The EU regulation on IAS will not exist anymore, but some aspects of the plant and animal health regulation will be still operational at national levels. Many ecosystems would be managed more actively, e.g. forests would be managed to maximize wood production, including the planting of species that increase yield value (including IAS). Freshwater invasive species are left unmanaged, and invasive species from the aquaculture pathways are being introduced in freshwater systems as food resources. Likewise, marine systems are highly influenced by overharvesting and the spread of marine IAS. Invasion literacy and awareness are low, and there is hardly any research dedicated to IAS impacts on biodiversity or how to mitigate for those.

While colonisation pressure (the number of species introduced) might decrease due to lower global trade compared to other trajectories, it would still be on the rise compared to today. Invasion debts would be an issue in the first 20 years. It is similar for propagule pressure, but while some alien species will have a very high propagule pressure compared to today (e.g. commercial IAS), some others will be actively controlled and managed. The latter is particularly true for alien species that are detrimental to human health, forestry or agriculture which will be strongly managed, although efficiency of measures is hampered by a lack of international collaboration and low scientific literacy and innovation.

In general, invasive species impacts and spread would exacerbate, favoured and in conjunction with other environmental pressures such as land use and climate changes. Management of biological invasions will not be a priority, and the spread and impact of invasions in natural areas will increase. Although IAS would more be passengers of environmental change, their impact on biodiversity would still be considerable as they would be largely left unmanaged. As societies become increasingly stressed with increasing climate change impacts, and international cooperation is limited, the management of alien species not directly threatening human livelihoods is increasingly becoming less relevant, favouring many alien species to expand.

**SCENARIO Eur-ASN B: BIG TECH RULES EUROPE**

| | |
|---|---|
| **Scenario breakout group** | Belinda Gallardo, Piero Genovesi, Marina Golivets, Chunlong Liu, Cristian Pérez-Granados (facilitator), Garry Peterson (facilitator), Jörg Priess, Peter Robertson, Wojciech Solarz |
| **Scenario key features** | **Keywords:** economic power, increasing trade, European homogenization, megacities, land abandonment, rural depopulation, disconnection from nature, increased land use, environmental degradation, loss of local knowledge, lack of monitoring, ecological surprise<br>**IAS specifics:** low regulations and coordination among countries to reduce IAS, large companies focus on rules to promote easy movement and trade (with little regard for nature or species), high propagule pressure, large pool of invaders (due to global trade), limited control and management of invasive species, urban novel ecosystems - many IAS species, invasive species in rural areas difficult to control due to depopulation, shifting baseline of what species expected |
| **Scenario downscaled from** | Global-ASN S11 'Globalized Corporation Society' (Scenarios Cluster B, see Roura-Pascual et al. 2021) served as the basis for downscaling to the European level |

### SCENARIO SUMMARY

The recession of the 2020s and the stimulus response to bail out companies lead to a distrust in governments due to their failure to stop the crisis and to a desire of people to look after their own. This leads to companies having increased power over European policy and a lack of mechanisms to control them (and their ability to hide in tax havens and behind financial instruments) because they are multinational and difficult to halt, sanction, or regulate. People have economic power but are economically stressed and increasingly disconnected from the environment. Most Europeans focus on urban life and improving their situation in the city. The response to the crisis of the 2020s accelerates land abandonment and rural depopulation. Rural depopulation enables the widespread expansion of intensive and commercial farming for markets, often done by large companies. Citizens show little interest in nature, ecosystem services or biodiversity and have little knowledge about ecology and IAS. Substantial trade without biosecurity leads to an exponential increase in IAS. There is a large increase in IAS propagule pressure (due to intentional introductions and novel pathways of unintentional introductions) and a decrease in coordinated management. This leads to a further homogenization of IAS in urban areas and the appearance of substantial numbers of IAS in rural areas. Rural depopulation leads to a lack of knowledge and control of IAS in rural areas; in combination with increasing climate change this leads to disturbed ecosystems, in which IAS are difficult to control. European regulations to prevent the introduction of IAS exist, but the public sector has little power, thus most management actions to control IAS are implemented by companies. Control of IAS is focused on economically damaging IAS, but is often slow, scarcely coordinated and ineffective due to fragmented responses and lack of deeper knowledge due to limited collaboration and knowledge exchange. Trade and river transport increase the spread of IAS in freshwater and marine ecosystems.

### IN DETAIL: WHAT DOES THIS SCENARIO IMPLY FOR EUROPE UP TO 2050?

#### REGARDING BROAD, CONTEXTUAL DEVELOPMENTS

**1. Political and institutional developments**

World economy will be dominated by US and Chinese-based companies (Microsoft, Google, Tencent...), but the EU and few EU companies are still among the global key-players setting the rules owing to their big contribution to the world economy. Brussels and Geneva also play a key role in global jurisdiction. The EU may have less power under this scenario due to different regulations that could be implemented in other areas, such as the US and China. Regulations implemented are not very useful and are controlled by big companies, with economy driving decision-making. It may partly fragmentate the EU with some countries creating different blocks, which may give place to inequality in wealth and access to education. However, a general homogenization is expected among areas of the EU, with increasing differences between rural and urban areas. The control and management of IAS is not an issue at EU scale, despite increasing propagule pressure under this scenario. However, at national scale various policies might be implemented to slow or halt the economic losses and increasing health risks produced by IAS, especially because the EU still has the capacity to develop new technology.

**2. Socio-economic and demographic developments**

EU population is slowly decreasing due to continued aging and low birth rate. The EU is still a politically and economically region that is attractive for immigrants. International migration may partly solve the declining population size. There is economic disparity and differences in wealth between rural and urban areas, with a general increase in farming. Environmental protection is no longer an issue, leading to the loss of ecosystem functions and favouring cascading effects (i.e. successful invasive species facilitate invasion of new species) of species invasions due to changed ecosystems.

**3. Culture, norms and values**

There is an increasing disconnection from nature in cities, with loss of knowledge of rural landscapes and ecosystems. This loss of knowledge leads to decreased focus on environmental changes and invasive species. People prefer to live in big cities, leading to highly invaded urban areas and abandoned landscapes. However, there is a large movement of goods, creating new problems for species invasions both in terms of propagules, establishment and spread. There is a different culture between rural and urban areas, and among countries depending on their economic situation. There is also a decreasing cooperation among countries, with some of them favoring economic isolation.

**4. Technological developments, science**

The development of technologies is largely controlled by the big corporations, with low competition among countries and reduced funding distributed to the scientific sector. There is little technological development oriented to managing ecosystems, landscapes and public goods; such as the spread of invasive species. Likewise, there is a lack of control and monitoring technologies and few methods are developed for managing invasions. Attention is paid just to those species with health impacts. In the long term

(2040-2050), substantial efforts might be made to reconnect school kids and citizens with nature, due to ever increasing environmental degradation and the growing number and impact of invasive species. An increase in climate adaptation and climate mitigation projects that involve green infrastructure and novel ecosystems begins to provide a novel pathway of new IAS into Europe, focussed on cities.

**5. Natural resources, ecological developments**

All across the EU countries and regions, land use is intensified. Wherever sufficient workforce is available, the growing amount of degraded lands and declining natural resources are used for bioenergy/biomaterial production. Many of the degraded and ecologically unstable lands suffer from biological invasions, causing the EU to lift the ban on glyphosate and other toxic agrochemicals to enable farmers, land owners and companies to continue agricultural production and (re-)establish bioenergy/biomaterial crops. Globalization gives place to a high propagule pressure with invasive species filling environmental niches across the globe. Environmental protection is not a concern, with pollution and climate change increasing. Depopulation of rural landscapes in Europe leads to more intensive and more extensive large-scale agriculture - focussed on high value and low value crops for biomaterials and biofuels. These areas may be growing new crops. This might be conversion of small-scale agriculture in central & eastern Europe, while land abandonment would likely continue in mountain regions of Europe. Bigger urban areas create big ecological problems, with a high consumption of resources, land use and materials. These changed ecosystems open the door for the establishment of many invasive species, and changes the opportunities between rural and urban areas, as well as among mountainous and coastal regions.

### REGARDING SPECIFIC IAS DYNAMICS AND MANAGEMENT

There is a high propagule pressure and a big pool of invaders due to globalized trade together with a reduction of biocontrol. Central countries, with a larger number of connections, would be more exposed than peripheral countries to biological invasions. This applies for both movement between river basins as well as trans-continental. Marine and aquatic invaders could be a bigger problem, since they suppose a lower threat to agriculture and might thus be even less likely to have a reactive response.

Movement of people to cities, would increase propagule pressure of terrestrial species in large urban areas. Urban areas may act as a shelter for invasive species, since they have different microhabitats, food nutrients and warmth from where invasive species may move to rural hinterlands through trade and transportation. Rural areas could become an invasion hotspot due to abandoned land use and rural depopulation, which may reduce their control and early detection. Agricultural intensification increases the probability of establishment and introducing alien species, with private farmers lacking funding to control invasive species.

There is a low coordination about management interventions of invasive species in Europe. There are European regulations to avoid the introduction of IAS, but the public sector has little power. Companies are not generally interested in the management and control of invasive species. Management of IAS is focussed on IAS that impact the activities of large companies. Companies and governments collaborate to monitor and control these species. The effect of these efforts will be reactive and focussed on impacts on agri-business. However, the effectiveness of implemented interventions will be low due to lack of coordination, lack of knowledge and lack of integrated approaches. These efforts are often successful in reacting to control new damaging IAS, but there is little action to prevent the arrival of new potentially damaging IAS for even these sectors.

### SCENARIO Eur-ASN C: GREEN LOCAL GOVERNANCE

| | |
|---|---|
| **Scenario breakout group** | Rubén Bernardo, Spyridon Flevaris, Marcus Hall, Petra Kutlesa, Guillaume Latombe, Teresa Pastor, Wolfgang Rabitsch, Núria Roura-Pascual (facilitator) |
| **Scenario key features** | **Keywords:** disintegration of the EU, development of regionalism, cooperation between European countries for certain concerns, participatory democracy, post-consumerism, degrowth, return to rural areas, environmental awareness, limited global understanding, low technological development, increased land sharing, increased fragmentation, limited efficiency of conservation efforts, limited climate change increase<br>**IAS specifics:** loss of EU regulations due to EU disintegration, reduced propagule pressure from outside Europe, secondary spreads within Europe difficult to manage, less efficient biosecurity because of lack of coordination, weakest link influence, less IAS but more widespread across Europe |
| **Scenario downscaled from** | Global-ASN S33 'Fairy Tale' (Scenarios Cluster C, see Roura-Pascual et al. 2021) served as the basis for downscaling to the European level |

### SCENARIO SUMMARY

By 2050, the EU as we know it (i.e. the common market promoting the exchange of goods, capital, services and labour) does not exist anymore and Europe sees the development of regionalism. It is not an increase in national populism and xenophobia, but a valorisation of local cultures and participatory democracy. Regional governments acquire greater influence because of the strong bottom-up participatory society, although there is still good cooperation among countries on certain political decisions or concerns (such as human health). Despite certain cooperation at a continental scale, there is a more superficial understanding of global (environmental) issues than currently. European society follows the degrowth paradigm, with less technological developments, and with the production of essential goods and services and consumption of local products. Eco-efficient, locally based production techniques are valued. People move from urban to rural areas. All these actions result in a reduction of GHG emissions and therefore a reduction of the impact of climate change. Remote working and remote learning is necessary and made easily accessible due to more distributed populations. Habitat fragmentation and increased land sharing are major pressures on the environment and the countryside, but mitigation measures, such as the creation of green corridors, are implemented. Nonetheless, some conservation efforts are less efficient due to the lack of a common environmental strategy. Because of isolation and reduced trade, the rate of introduction of new alien species coming from outside Europe decreases. However, the further spread of already established IAS is difficult to manage because of less efficient biosecurity measures, not coordinated at a continental scale.

### IN DETAIL: WHAT DOES THIS SCENARIO IMPLY FOR EUROPE UP TO 2050?

#### REGARDING BROAD, CONTEXTUAL DEVELOPMENTS

**1. Political and institutional developments**

This scenario sees the development of regionalism, i.e. of more local structures. It is not an increase in national populism and xenophobia, but a valorisation of local cultures and participatory democracy. Top-down structures do not progress because of the strong bottom-up participatory society. There is an acknowledgement and respect of other cultures without merging of these cultures. Democratic and human-right principles are highly valued.

The EU as we know it (i.e. the common market promoting the exchange of goods, capital, services and labour) does not exist anymore, but there is some coordination structure amongst countries, which becomes the main “consultant” organ among European states. This would take the form of a European equivalent to the UN and sub-organisations (such as the UNESCO and UNEP). There would also be organisations functioning in similar ways as the WHO. States do not need to follow its recommendations, but most of them do because they “feel” pressured to do it (to comply with global mandates).

There is no overarching enforcement. Instead, self-determination of countries and regions is promoted. Nonetheless, there is a general understanding that the fate of a country depends in part on the actions of other countries, and the legacy of the EU promotes good cooperation through participatory communications for issues that affect all countries, including IAS (e.g. UK-UE after Brexit). This allows avoiding a tragedy of the commons and is inspired by Ostrom's eight design principles for the management of common-pool resources, particularly principles (1) Commons need to have clearly defined boundaries; (2) Rules should fit local circumstances; and (8) Commons work best when nested within larger networks (Ostrom, 1990).

**2. Socio-economic and demographic developments**

Countries follow a degrowth paradigm, with less technological developments and more emphasis on the local production of essential goods and services (food, health, etc.). Although there is an emphasis on the consumption of local products, each country is not completely independent, and some level of cooperation (preferably among neighbouring countries) enables to fill the gaps. Because of self-determination (understood as an international principle), there is a variety of relationships with other countries outside Europe (e.g. Eastern countries tend to exchange goods with Russia, whereas the UK is more connected to the US). It may lead to more egalitarian societies at the regional level (since the regionalization can also be within the countries), but more divergences between EU countries.

There is a movement of people from urban to rural areas, with the consequent redistribution of resources/services (e.g. schools, hospitals, transport network, etc.) across geographic regions. Economic investments to adjust to these changes will reduce or alter the budget available for other aspects, such as the management of IAS or research programs. The absence of centralised structures makes it harder to get some services (such as health care). Remote working and learning is easily possible (and necessary) due to a more distributed population. No huge changes in demographic rate.

**3. Culture, norms and values**

Local culture and ecology are promoted through education. Strong local cultural values and a strong care for the environment will tend to promote the control of IAS to preserve native ecosystems. Despite cooperation between European states, there is a more naïve perspective regarding global issues, and a limited sense of global understanding, which might be problematic when dealing with environmental (global) issues. However, there is more transparency and good participation from society that can (partly) counteract the effects of divergences between countries.

Native species will be valued and therefore IAS will be managed. However, there will be animalist groups against the eradication of IAS animals and other groups favourable to keep some emblematic IAS species/patches, but they will do their best to keep IAS plants and animals under control. Given the biodiversity in Europe, nature-based solutions are likely to be implementable with the native pool. Overall, the final number of IAS will not further rise.

**4. Technological developments, science**

At the country level, the redistribution of resources to meet the demands of the growing rural population will reduce the financing for technological developments and science. At continental level, the disappearance of UE institutions that promoted the collaboration between different countries will cause more regional hubs of innovation and less development at a global scale. Consequently developing international big projects such as space race (e.g. satellites) will be more challenging and technological developments will be mostly directed at food production, health systems, and primary needs, but with the limitations mentioned above (leading to slowing innovation in production sectors, fewer patents, and less medical advancement). Nature-based solutions will be preferred with a valorisation of eco-efficient and traditional approaches that were used in the past.

**5. Natural resources, ecological developments**

There would be an increased land sharing, due to increased rural population. It would be a major pressure on the environment/countryside because there would be a need to increase regional connections (leading to fragmentation). However, the new green mind may also facilitate the investment in mitigation measures (e.g. the creation of green corridors) to reduce habitat fragmentation. Land sparring will occur when possible, but will likely be limited due to lack of space. Rural standards and local production will go up, as well as an ecologically sensitive management of urban, periurban, and rural systems. The local application of Ostrom’s eight design principles will lead to sustainable use of local natural resources.

However, nature does not follow political borders. Commons work best when nested within larger networks. Some things can be managed locally, but some others might need wider regional cooperation – for example, an irrigation network might depend on a river that others also draw on upstream. So, under this scenario, the conservation efforts for large conservation or similar areas are less effective and sometimes impossible due to regionalisation. With the lesser perception of the need for conservation at the global level, regions that may have high conservation value may be more threatened by being subjected to the management of regional governments that may have a different perception of the value of some habitats or ecosystems. In general, as happened with science, it would be difficult to perform global conservation projects and assessments.

We assume that the rest of the world follows a similar "green local governance" scenario. There is a global reduction in the increment of greenhouse gas emissions, which slows the pace of climate change. Global warming is limited to ~2º Celsius degrees compared to pre-industrial levels in 2050s.

### REGARDING SPECIFIC IAS DYNAMICS AND MANAGEMENT

Because of isolation and incentives to implement prevention measures, introduction rates of IAS coming from outside Europe would be reduced. Also, because climate change will be slowed down, and there will be a reduced chance of having favourable environments for novel IAS. But once IAS are established in Europe, natural secondary spreads will be difficult to manage. Spread will nonetheless occur in a more diffused manner, with less long-distance dispersal within Europe due to decreased commercial exchanges and movements of people. There will be cooperation once the problem has been identified, but less efficient biosecurity because it is not coordinated at a continental scale (because of different country's capacities, interests, beliefs, priorities, etc).

Finally, countries do not have the same understanding of nature resulting in different species being targeted by local management measures, leading to increased risks of IAS reaching new countries. Additionally, because of regionalism, management priorities across countries are also divergent and priorities at local scales can be different from the country ones. In a nutshell, the disintegration of the EU will reduce biosecurity capacity at European level, all efforts by individual countries (or regions or stakeholders) will be undermined by the weakest link. Although trade will be reduced and the rate of introduction of new alien species decreases, the management of IAS will still be challenging. Overall there would be fewer IAS, but more widespread across Europe.

**SCENARIO Eur-ASN D: TECHNOLOGICAL (PSEUDO-)PANACEA**

| | |
|---|---|
| **Scenario breakout group** | David Aldridge, Sven Bacher, Lluís Brotons, Bernd Lenzner, Konrad Pagitz, Wolf-Christian Saul (facilitator), Rob Tanner, Montserrat Vilà |
| **Scenario key features** | **Keywords:** cooperative Europe, effective biosecurity, "green" but urban lifestyle, high trade volume, technological advancement, urbano-technophile society<br>**IAS specifics:** effective biosecurity through extended legislation and strong cooperation between European countries, but also intensified border controls; high trade volume means high propagule pressure; IAS predominantly in cities, semi-urban areas, transportation networks outside cities, areas of intensive agriculture (i.e. highly technological, artificial, novel ecosystems); applied research strongly promoted: advanced technology for prevention, detection, monitoring, control; interoperability of IAS databases across countries; positive effects of IAS taken into account |
| **Scenario downscaled from** | Global-ASN S12 'Hipster/Techno Society' (Scenarios Cluster D in Roura-Pascual et al. 2021) served as the basis for downscaling to the European level |

### SCENARIO SUMMARY

European nations in this scenario cooperate strongly, with fast technological advancement, large trade volumes and high biosecurity being the prime societal and policy objectives. Throughout Europe (beyond EU borders), agencies responsible for environmental policy are significantly strengthened and biosecurity legislation is strictly enforced. Despite strong cooperation, the reintroduction of intra-European border controls for biosecurity reasons leads to the end of free movement within the EU. European societies are highly urbanophile and concentrate in "Smart cities". The urban lifestyle is supported by intensive, industrialized agriculture in rural areas and results in a decoupling from nature. Citizens strongly believe that technological progress can solve all current and future problems. Education is equally accessible to all parts of society and the general public acknowledges environmental issues (e.g. the impact of IAS). There is substantial cooperation regarding conservation and biosecurity measures, but policies are reactive rather than proactive. The strict regulations entail a degree of rigidity in societal life that fosters the development of rather complacent, passive societies. To a limited extent, implementation of the regulations varies among countries due to differences in types of governance, cultural legacies and values. Europe, and its individual nations, are engaged in an all-engulfing race to stay ahead of potential problems and of competitors by developing new technological solutions. Technological research and entrepreneurship is strongly promoted. Technologies for reducing the ecological footprint of various activities are available and implemented across Europe. Societies have a high, but not increasing ecological footprint. Catastrophic events, such as fires and floods, are under control using the latest technology, and further biodiversity loss is halted. Despite intensive global trade, the rate of IAS establishment and spread is low because of strong and diligent biosecurity measures, efficient risk assessments and other precautionary measures. As a result, different sets of IAS occur in cities and semi-urban environments as well as transportation networks outside cities and areas of intensive agriculture (i.e. predominantly in highly technological and artificial novel ecosystems). IAS management is supported by technological advances in automated and remote data collection with very high coverage, at large spatial and temporal scales and using standardized protocols in Europe.

### IN DETAIL: WHAT DOES THIS SCENARIO IMPLY FOR EUROPE UP TO 2050?

#### REGARDING BROAD, CONTEXTUAL DEVELOPMENTS

**1. Political and institutional developments**

Overall, European nations in this scenario cooperate strongly, with fast technological advancement, large trade volumes and high biosecurity being the prime societal and policy objectives. To ensure highly effective biosecurity on a European level, EU agencies and agencies of non-Member states responsible for environmental policy are significantly strengthened (e.g. in terms of their funding and weight in the political decision process). Further, existing biosecurity legislations (e.g. the Ballast water convention) as well as species lists for regulating IAS have been extended and strictly enforced throughout Europe. These are based on more advanced, standardized and comprehensive evidence-based cost-benefit analyses of IAS, which also recognize the intangible, non-monetizable costs of IAS having an impact on biodiversity beyond economically important crops. There is a functioning collaboration structure between European countries (EU member and non-member states) to promote good coordination and more equity in the countries' responses to IAS (concerted action with more regional than national focus). In particular, collaboration between EU agencies responsible for environment, trade and agriculture is strengthened, while plant health and the environmental sector have strong links to ensure rapid and knowledgeable biosecurity responses.

Europe has a long history of multi-country agreements, and the effectiveness of their implementation is relatively high compared to other parts of the world. However, despite the strong cooperation, Europe is still made up of individual nations, and national interests still play a role. For instance, subliminal friction originates from the fact that different types of societies and governance co-exist within Europe, ranging from societies with a strong overall top-down governance to more permissive societies with strong reliance on market-driven (bottom-up) self-regulation. Differences between these influence the respective way of conceiving and implementing regulations and technological solutions as well as their consequences. To a limited extent, thus, implementation of the regulations varies among countries due to differences in types of governance, cultural legacies and values (e.g. due to some countries or industrial branches retarding the implementation of certain regulations and technological advances when it serves their interests). Furthermore, many new legislative biosecurity instruments are developed which also entail more border controls for products and people within EU member countries (and a substantial increase in bureaucracy). This leads to the abolishment of the Schengen agreement.

Key trade-off in this context: more biosecurity (e.g. border interceptions) vs. more free trade.

**2. Socio-economic and demographic developments**

Economic equality within Europe is high. Pro-trade agreements are in place (for within the EU, for Europe and globally), with trade increasing due to global e-commerce and new trade technologies. The technological infrastructure is highly developed across

European national borders and in rural areas, promoting the strong development of e-commerce (on-demand delivery etc). As a consequence, the diversity, dispersion and volume of traded goods is high, as is propagule pressure of IAS. Environmental and IAS regulations are effectively enforced in the economic sector: enterprises by default implement environment-friendly and sustainable processes for production and distribution (e.g. low carbon and land-use footprint; guaranteed by innovative technology), as well as measures against the spread of IAS that could occur through their activities. European societies are attracted to living in large cities, having strong affinities to a green but urban lifestyle with amenities such as development of green infrastructure, fast and widespread public transportation, affordable accommodation etc. This lifestyle is supported by intensive, industrialized agriculture in rural areas. Access to health care is relatively egalitarian, leading to long life expectancies across society as well as a low population growth[1] . Targeted technological development addresses the needs of all generations in society, including of the elderly people for whom it might otherwise be difficult to keep up with the technological advancements.

Europe (and its individual nations) are engaged in an all-engulfing, exhaustive race to stay ahead of potential problems and of competitors by developing new technological solutions. This comes with drawbacks related to e.g. at least temporary unsustainable use of resources and funding shortage of other societal sectors/activities. It may also prompt (semi-)illegal actions from some actors to gain advantages, e.g. actively seeking and exploiting loopholes in regulations or manipulating data to comply with regulations. Such race may be more pronounced in those European nations at the permissive, market-driven end of the societal spectrum than in more top-down regulated societies. The latter, on the other hand, may experience difficulties because top-down planning is less efficient in recognizing upcoming problems and finding innovative solutions due to missing incentives for people to develop them (no personal gain).

**3. Culture, norms and values**

The highly urbanophile lifestyle of European societies results in a decoupling from nature, resulting in a skewed perception of wilderness and a decline in taxonomic knowledge. There is a strong belief that technological progress can solve all current and future problems. Education is equally accessible to all parts of society, based on high educational standards with heavy emphasis on science and technology (MINT courses). As a result, and despite the decoupling from nature in daily life, the general public is knowledgeable about ecological concepts and acknowledges environmental issues (e.g. the impact of IAS). It accepts the need for conservation and biosecurity measures, being cooperative but not necessarily proactive. In general, different cultural legacies and values across Europe still have some impact on the effectiveness of IAS management. For instance, despite awareness in the general public of possible negative impacts of IAS, concerns against IAS control measures are not completely dispelled even by the availability of methods making use of advanced technology. "Eco-"products (e.g. local products, slow food) are generally favoured even if it means paying more, but short-term trends and (lifestyle) fashions for products also open new trade routes (regional, global), increase the influx (frequency and volume) of novel IAS and represent a constant challenge to biosecurity. Positive effects of IAS are also getting more attention, particularly those related to novel ecosystems emerging in urban areas and in the context of natural capital (e.g. monetary value or other benefits counterbalance concerns about IAS).

European societies focus on the potential of technological solutions to such extent that they neglect other important cultural aspects of human and non-human life. Life becomes more impersonal as jobs become asocial, and personal care, face-to face discussions, socializing and a vibrant cultural sector are of low importance. In schools and universities, the strong focus on MINT courses comes at the expense of education in the humanities (nevertheless, artistic and literary expressions about IAS and other environmental topics do exist due to the high public awareness). Overreliance on big data for explaining all phenomena neglects the influence of people's attitudes and emotions (Dataism). Further, the strict regulations to ensure high biosecurity alongside fast technological advancement and large trade volumes entail a degree of rigidity/inflexibility in societal life that fosters the development of rather complacent, passive societies. Ethical concerns become of subordinate importance in Europe (e.g. privacy and personal rights: tracking of people is acceptable to prevent spread of IAS). Depending on the society type (ranging from relatively permissive to strongly top-down regulated), this may prompt discord or even resistance ("subversive action") at least in (smaller) parts of society that strive to maintain their personal freedom and privacy.

Key trade-off in this context: buying local, organic/environmentally friendly products vs. indulging in exotic products/fashions/subculture (incl. travel habits)

**4. Technological developments, science**

Technological research and entrepreneurship is strongly promoted at the European level (e.g. high funding, tax incentives, collaborative research programmes). Besides increased research related to trade and transport innovations, research on IAS is applied and technical, strongly geared towards problem-solving and predictions based on modeling and simulations. By contrast, funding for basic research (incl. taxonomy and natural history of IAS) is low. Technological advances have been achieved in relation to IAS-detection via remote sensing (e.g. using drones), high-performance screening technology at points of entry (e.g. harbors, airports, logistics centres), eDNA and other identification techniques, as well as breeding of sterile plants for horticulture or sterilized animals for pet trade. Citizen science approaches are widely applied and supported by cheap and easy-to-use technology (e.g. apps for entertaining forms of communicating the IAS issue rather than actively producing data with the help of citizen). As a major step towards effective, coordinated biosecurity in Europe, the interoperability and harmonization of relevant databases across countries has been increased. Further-reaching technological advances have been made in automated and remote data collection with very high coverage, at large spatial and temporal scales and using standardized protocols in Europe. This includes, for instance, all kinds of sensors, cameras, eDNA sampling (e.g. ballast water), and AI algorithms for species recognition. There is high usage of automated culturomics approaches (i.e. scraping information from platforms like Google, Instagram, etc.) that identify species in all kinds of resources, including even those that do not specifically target IAS (e.g. holiday pictures). Cities are “Smart Cities” using information and communication technology to optimize the efficiency of city operations and services and connect to citizens. All this entails a strong increment in big data and the need for automated data analysis. Data and analyses from automated and algorithm-based data collection technologies are subject to strict quality checks/assessments. Securing assessment reliability, however, is a constant challenge and connected with high costs due to the lack of enough personnel trained in taxonomy and natural history (of IAS). Overall, there are technological solutions in place for dealing with climate change (climate engineering, renewable energy production, low-carbon technology) and land-use change (e.g. advancing use of existing agricultural land, aquaculture, GMOs). Technologies for mitigating the ecological footprint at larger scales are available and implemented across Europe, and catastrophic events (fires, floods) are under technological control. More environmentally friendly and targeted chemical products for controlling IAS of major concern are available or replaced by other management methods, e.g. genetic control techniques. The large economic and educational capacity of European countries allows a more effective implementation of the measures foreseen by the scenario (technological solutions in general and targeting IAS) compared to other parts of the world.

Key trade-offs in this context: applied vs. basic research; data mining vs. natural history studies; innovative technology entailing new biosecurity challenges, e.g. novel organisms (GMO, etc.).

**5. Natural resources, ecological developments**

Technologies for mitigating the ecological footprint at larger scales are available and implemented across Europe. Societies have a high, but stabilized ecological footprint. Cities are focal points in European societies' existence and across Europe efforts have been successful to transform them into "Smart Cities" that are more efficient, more sustainable, healthier, cleaner and more liveable. In order to protect natural resources, value and supply chains are optimized for sustainability (e.g. circular economy, more efficient transportation). Temporary unsustainable use of resources results from the technological race in which Europe takes part. There is also intensive exploration of opportunities for natural "medication". Strong land-sparing land use is in place, a result of such tendencies in Europe's recent history. Which pristine areas are to be maintained under protection for recreation or for delivering ecosystem services is decided on the European level. This also includes increased experimentation regarding the transformation of natural systems into functional systems that deliver specific services. With the help of advanced technology, exploited and unproductive areas are put to some kind of use within short time so that only few neglected areas exist in which nature could reclaim territory by itself. Overall, further biodiversity loss is halted and catastrophic events (fires, floods) are under technological control.

### REGARDING SPECIFIC IAS DYNAMICS AND MANAGEMENT

The number of introduced IAS is relatively high because of intensive global trade and transport, but countered by highly effective pre-border biosecurity efforts. The rate of IAS establishment is low, because of strong and diligent biosecurity measures, efficient risk assessments and other precautionary measures. Different sets of IAS occur in cities and semi-urban environments as well as transportation networks outside the cities and areas of intensive agriculture (i.e. predominantly in highly technological and artificial novel ecosystems). For instance, urban areas may have higher proportions of IAS that provide ecosystem services for urban life (e.g. related to aesthetics, shading etc.) than rural areas where more IAS related to (and potentially impacting) production systems may be present. The secondary spread outside cities occurs as a result of decentralized commodity distribution across Europe (using e.g. drones and self-driving cars) as well as recreation activities following the city–rural divide. However, it is limited in regard to protected areas due to high technological response capacities reserved for them. Facilitation of IAS through other drivers of global change is limited, but has not stopped. Further loss of the remaining biodiversity is halted, as is also the loss of ecosystem services, which are sometimes artificially enhanced. Similarly, the number of IAS having negative impacts on plant, animal and human health is low due to high technological response capacities.

IAS management mainly relies on applied and technical research, strongly geared towards problem-solving and forecasting based on algorithms and simulations. It is backed up by technological advances in automated and remote data collection with very high coverage, at large spatial and temporal scales and using standardized protocols in Europe. Relevant local, national, regional and European databases are harmonized and interoperable across countries. For the sake of high biosecurity, IAS target lists are compiled more based on facts and less subject to (economically driven) negotiations. Insofar, management is more evidence-based than ever before. Also positive effects of IAS are getting more attention, particularly those related to novel ecosystems emerging in urban areas and in the context of natural capital (which is where monetary considerations come in again). With fast technological advancement, large trade volumes and high biosecurity being prime societal and policy objectives, ethical concerns become of subordinate importance in Europe (e.g. privacy and personal rights: tracking of people is acceptable to prevent spread of IAS).

Key trade-off in this context: potential availability of advanced and effective biosecurity technology vs. high costs for their actual use (i.e. are we willing or able to pay)?
